# SERPINB5-TGF-β signalling modulates desmoplakin membrane localization and ameliorates pemphigus vulgaris skin blistering

**DOI:** 10.1101/2024.10.15.618475

**Authors:** Maitreyi Rathod, Aude Zimmermann, Henriette Franz, Tomás Cunha, Dario Didona, Michael Hertl, Enno Schmidt, Volker Spindler

## Abstract

Impairment of desmosomal cell-cell adhesion leads to several life-threatening diseases such as the autoimmune skin blistering disorder pemphigus vulgaris (PV). Disease management strategies that stabilize intercellular adhesion, in addition to the existing immunosuppression therapies, may result in improved clinical outcomes. Previous findings showed that the serine protease inhibitor SERPINB5 promotes intercellular adhesion by binding to and regulating the localization of the desmosomal adapter molecule desmoplakin (DSP) at the plasma membrane. We here show that SERPINB5 overexpression prevents PV-IgG-mediated loss of cell-cell adhesion and the loss of DSP from the cell membrane. We mechanistically demonstrate that SERPINB5 loss deregulates TGF-β signalling, a pathway known to destabilize DSP in keratinocytes. TGF-β signalling was also activated in skin biopsies of PV patients and keratinocytes treated with PV autoantibodies, suggesting a contribution to disease. Inhibition of TGF-β activation ameliorated PV-IgG-mediated loss of cell-cell adhesion, increased DSP membrane expression, and prevented PV-IgG-induced blister formation in a human *ex-vivo* skin model. Together, SERPINB5 modulates DSP and intercellular adhesion through the regulation of TGF-β signalling. Further, TGF-β signalling was identified as a potential target for pemphigus treatment.

## Introduction

The skin provides a barrier against external insults and is critical for the homeostasis of the organism. Keratinocytes represent the major cell type within the epidermis, the epithelium of the skin. The epidermis is exposed to constantly changing mechanical stresses and its integrity relies on strong connections of keratinocytes, which provide the necessary resilience against the acting forces^1^. Desmosomes, due to their molecular arrangement and linkage to the intermediate filament (IF) cytoskeleton, are ideally suited to facilitate these robust intercellular adhesions^2,3^. Accordingly, desmosomes are abundant in tissues which are exposed to high mechanical forces such as the epidermis or the myocardium^4^. Desmosomes are made up of an intercellular core of the desmosomal cadherins desmoglein (DSG) 1-4 and desmocollin (DSC) 1-3. These cadherins are coupled to IFs through intracellular adaptor proteins such as plakoglobin, plakophilins (PKP) and desmoplakin (DSP).^3^

Dysfunction in desmosomal cell-cell adhesion leads to severe pathologies^5^. Mutations of desmosomal proteins are directly associated with arrhythmogenic cardiomyopathy^6,7^. Further, autoantibodies targeting DSG1 and or DSG3 lead to the potentially fatal autoimmune skin blistering diseases pemphigus vulgaris (PV) and pemphigus foliaceus (PF)^8–10^. The microscopic hallmarks of pemphigus are intraepithelial blister formation and acantholysis, which is the loss of adhesion between keratinocytes. Mechanistically, it has been shown that autoantibodies induce steric hindrance, altered protein turnover, and aberrant signalling involving p38-MAPK, SRC, EGFR, and PKC, which all contribute to the disease outcome^11^. Current treatment strategies for PV rely on immune suppression strategies such as corticosteroid administration, steroids-sparing immunosuppressive drugs (e.g. azathioprine and mycophenolate mofetil) and B-cell depletion ^12,13^, which are associated with profound side-effects. Given the causal relationships between dysfunctional desmosomal adhesion and clinical phenotype, strategies which stabilize intercellular adhesion may serve as targeted therapeutic options. Thus, the exploitation of known mechanisms regulating membrane trafficking, expression and turnover of desmosomal molecules may serve an unmet clinical need.

In the current study, we show that SERPINB5 rescues autoantibody-mediated loss of cell cohesion through enhancing DSP expression at the cell membrane. SERPINB5 is a non-classical SERPIN, which is believed to not inhibit serine proteases^14^. SERPINB5, also called MASPIN, was first identified in breast cancer as a tumor suppressor. It was associated with migration and adhesion of cells^15^, however the underlying mechanisms are not well understood. It was previously shown that TGF-β signalling negatively modulates cell-cell adhesion and DSP expression in keratinocytes^16^ and TGF-β in cooperation with p53 positively modulates SERPINB5 expression in mammary epithelial cells^17^. We here identified TGF-β signalling as a mediator of cell-cell adhesion loss in PV patient samples. Inhibiting TGF-β activation prevented loss of intercellular adhesion *in vitro* and blister formation *ex-vivo* in human skin.

## Results

### SERPINB5 rescues PV-mediated loss of intercellular adhesion

Immunostaining-based analysis of DSP in SERPINB5 knockdown HaCaT keratinocytes (**Figure S1A**) confirmed our recent finding^18^ that SERPINB5 is a positive regulator of DSP membrane localization, while DSG3 localization was unaltered (**Figure 1A**). This is also in line with our results that SERPINB5 is required for normal intercellular adhesive strength in keratinocytes^18^. To study the effect of SERPINB5 on PV-mediated loss of cell cohesion, we overexpressed SERPINB5 in keratinocytes (**Figure S1B, C).** Dissociation assays showed that overexpression of SERPINB5 rescued loss of cell-cell adhesion in HaCaT and primary human keratinocytes (NHEK) induced by incubation with autoantibody fractions of PV patients (PV-IgG) (**Figure 1B, C**). These findings were confirmed using PX4_3, a single chain variable fragment (scFv) cloned from a PV patient B cell repertoire which targets both DSG3 and DSG1^19^ (**Figure 1D**).

**Figure 1.**
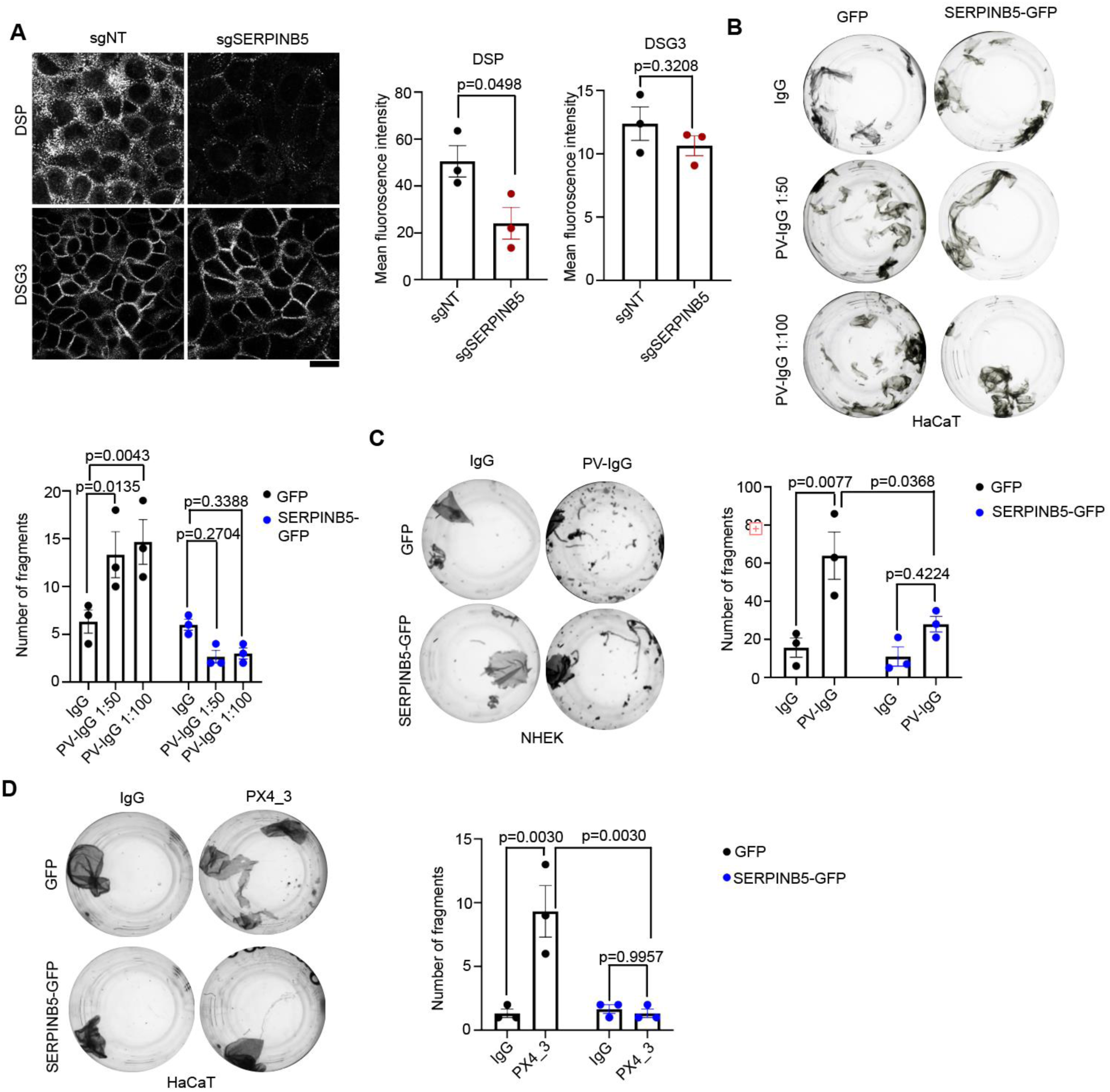
**A)** Immunofluorescence staining of sgNT1 and sgSERPINB5 HaCaT cells using DSG3 and DSP antibodies. Scale bar = 10 μm. Quantification of DSP and DSG3 mean fluorescence intensity of 3 independent experiments are shown. Each data point represents one biological replicate. Unpaired Students t-test used for statistical analysis. **B)** Dispase-based dissociation assay of HaCaT cells overexpressing GFP or SERPINB5-GFP and treated with IgG or PV-IgG at indicated concentrations. Representative images and quantifications of n=3 are shown. Two-way-ANOVA, SIDAK correction used for statistical analysis. **C)** Dispase-based dissociation assay of NHEK cells overexpressing GFP or SERPINB5-GFP and treated with IgG or PV-IgG (1:100). Representative images and quantifications of n=3 are shown. Two-way-ANOVA, Tukey’s multiple comparison used for statistical analysis. **D)** Dispase-based dissociation assay of HaCaT cells overexpressing GFP or SERPINB5-GFP and treated with IgG or PX4_3 (1:100). Representative images and quantifications of n=3 are shown. Two-way-ANOVA, Tukey’s multiple comparison used for statistical analysis.

### SERPINB5 stabilizes DSP at the cell membrane under PV-IgG conditions

Next, we analysed the localization of DSG3 in NHEK cells in the context of SERPINB5 overexpression. DSG3 staining showed the well-established fragmented pattern in the membrane and a general reduction at the cell surface upon treatment with PV-IgG, which was unaltered in the presence of SERPINB5-GFP (**Figure 2A**). However, staining of the central desmosome constituent DSP revealed that the reduction at the membrane upon treatment with PV-IgG was prevented by SERPINB5 overexpression (**Figure 2B**). This suggests that SERPINB5 stabilizes DSP localization independently of DSG3 under PV-IgG exposure and rescues loss of cell-cell adhesion.

**Figure 2.**
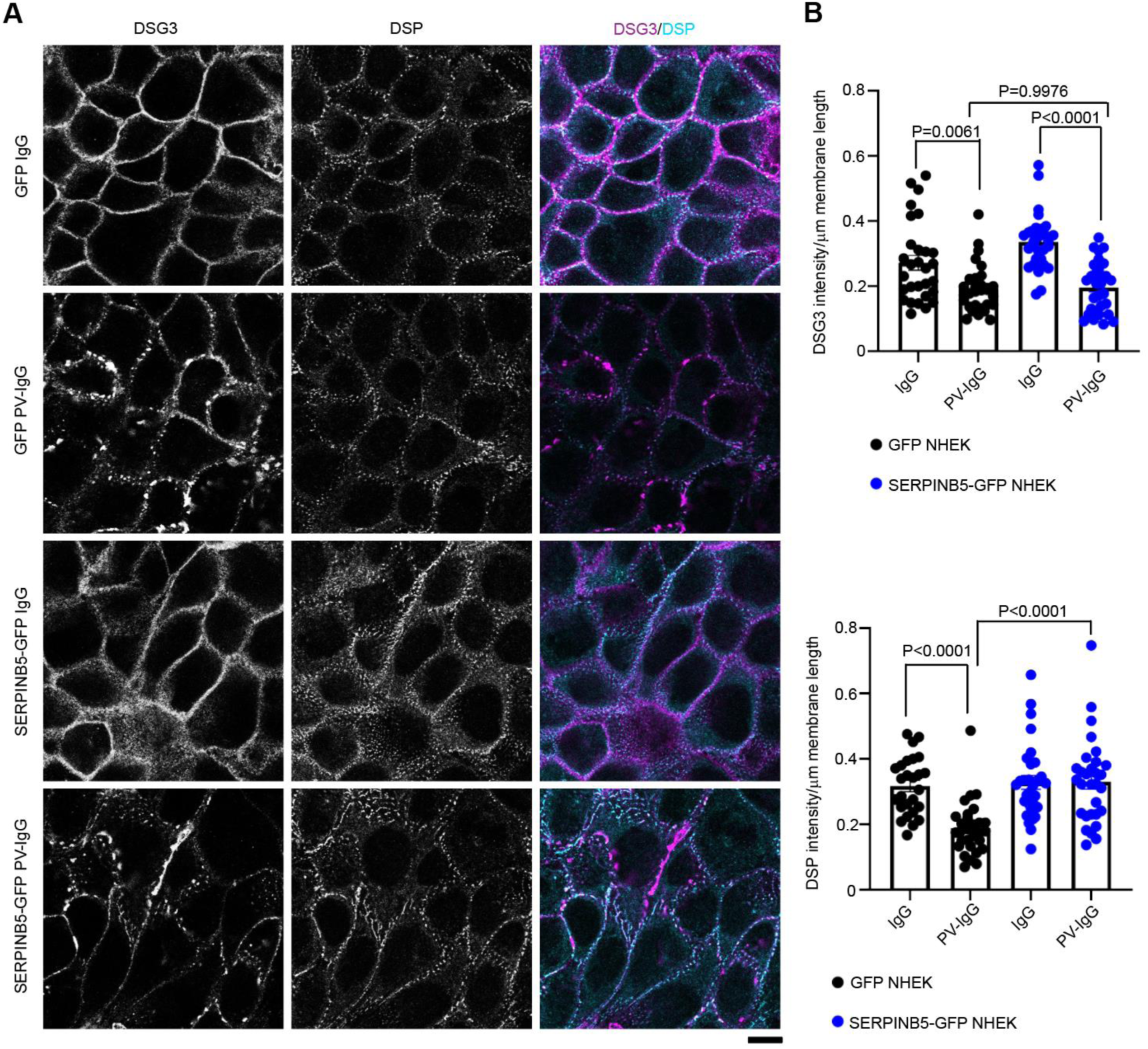
**A)** Immunofluorescence staining of DSG3 and DSP in NHEK cells expressing GFP or SERPINB5-GFP respectively. Scale bar = 10 μm. **B)** Quantification of DSG3 and DSP fluorescence intensity/membrane length of individual cells from 3 independent experiments are shown. One-way-ANOVA, Tukey’s multiple comparison used for statistical analysis.

### SERPINB5 regulates TGF-β signalling, which is activated in PV patient samples

TGF-β signalling is a known negative regulator of DSP expression and cell-cell adhesion^16^. Interestingly, knockdown (KD) of SERPINB5 in HaCaT keratinocytes led to significantly enhanced levels of pSMAD2/3 and pSMAD1/3/5 (**Figure 3A**). Inhibiting the activation of SMAD2/3 and SMAD1/3/5 by the small molecule inhibitor GW788388 (GW788) (**Figure S1D**), in SERPINB5 KD localization at the cell membrane (**Figure 3C**). Given that DSP localization at the membrane was impaired both in SERPINB5 knockdown cells and in keratinocytes incubated with PV-IgG, we evaluated changes in TGF-β signalling in PV patient biopsies. Indeed, analysis of nine healthy control and six PV patient samples revealed enhanced levels of pSMAD2/3 in PV epidermis (**Figure 3D**). Furthermore, keratinocytes treated with PV-IgG and PX4_3 showed elevated pSMAD2/3 and pSMAD1/3/5 (**Figure 3E**) levels, indicating that PV-IgG activates TGF-β signalling. Importantly, SERPINB5 levels were unaltered upon PV-IgG treatment (**Figure S1E**), suggesting that PV-IgG activates the TGF-β pathway independently of SERPINB5.

**Figure 3.**
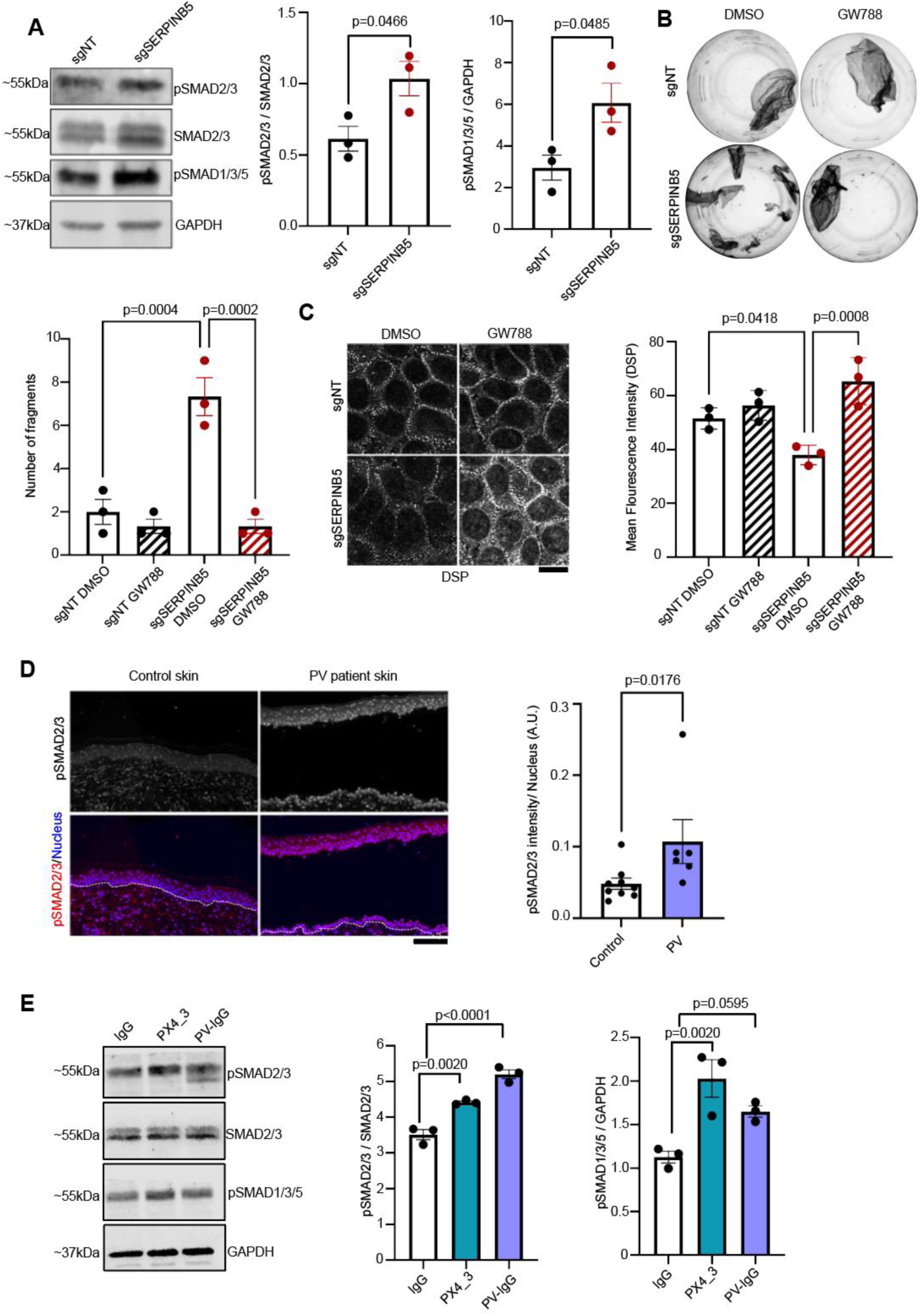
**A)** Western blot analysis of sgNT and sgSERPINB5 HaCaT cell lysates using pSMAD2/3, SMAD2/3, pSMAD1/3/5 and GAPDH antibodies. Representative Western blot images and quantifications of indicated proteins (n=3) are shown. Unpaired Students t-test was used for statistical analysis. **B)** Dispase-based dissociation assay of HaCaT cells with sgNT or sgSERPINB5, in combination with either DMSO or GW788388 (GW788) for 24 hours. Representative images and quantifications of n=3 are shown. One-way-ANOVA, SIDAK multiple comparison used for statistical analysis. **C)** Immunofluorescence staining of DSP in HaCaT cells expressing sgNT or sgSERPINB5 treated with DMSO or GW788. Scale bar = 10 μm. Quantification of DSP mean fluorescence intensity of 3 independent experiments are shown. Each data point represents one biological replicate. One-way-ANOVA, Tukey’s multiple comparison used for statistical analysis. **D)** Immunofluorescence staining of pSMAD2/3 in human epidermal biopsy sections from healthy controls or PV patients. DAPI served to visualize nuclei. Scale bar = 10 μm. Quantification shows the mean intensity of pSMAD2/3 per nucleus (n=9 controls and n=6 patients). Unpaired Students t-test used for statistical analysis. **E)** Western blot analysis of HaCaT lysates from cells incubated with IgG, PX4_3 or PV-IgG for 24 hours, using pSMAD2/3, SMAD2/3, pSMAD1/3/5 and GAPDH antibodies. Representative Western blot images and quantifications of indicated proteins (n=3) are shown. One-way-ANOVA, Dunnett’s correction used for statistical analysis.

### TGF-β inhibition ameliorates PV-IgG-mediated cell dissociation in cultured keratinocytes and acantholysis in an ex-vivo human skin model

Our findings suggest TGF-β signalling as a mechanism contributing to loss of cell-cell adhesion in PV. To address this, we inhibited the activation of SMAD2/3 and SMAD1/3/5 by the small molecule inhibitor GW788. Co-incubation of either HaCaT keratinocytes or NHEKs together with PV-IgG and GW788 prevented loss of cell-cell adhesion (**Figure 4A, B**). Similarly, PX4_3-mediated loss of cell-cell adhesion was blocked by GW788 treatment (**Figure 4C**). Further, inhibition of TGF-β signalling significantly increased DSP localization at the cell membrane in both IgG control and PV-IgG treated NHEK, suggesting that the inhibition of TGF-β signalling rescues loss of cell-cell adhesion through enhancing DSP membrane expression under PV-IgG influence, whereas DSG3 remains unchanged (**Figure 4D**).

**Figure 4.**
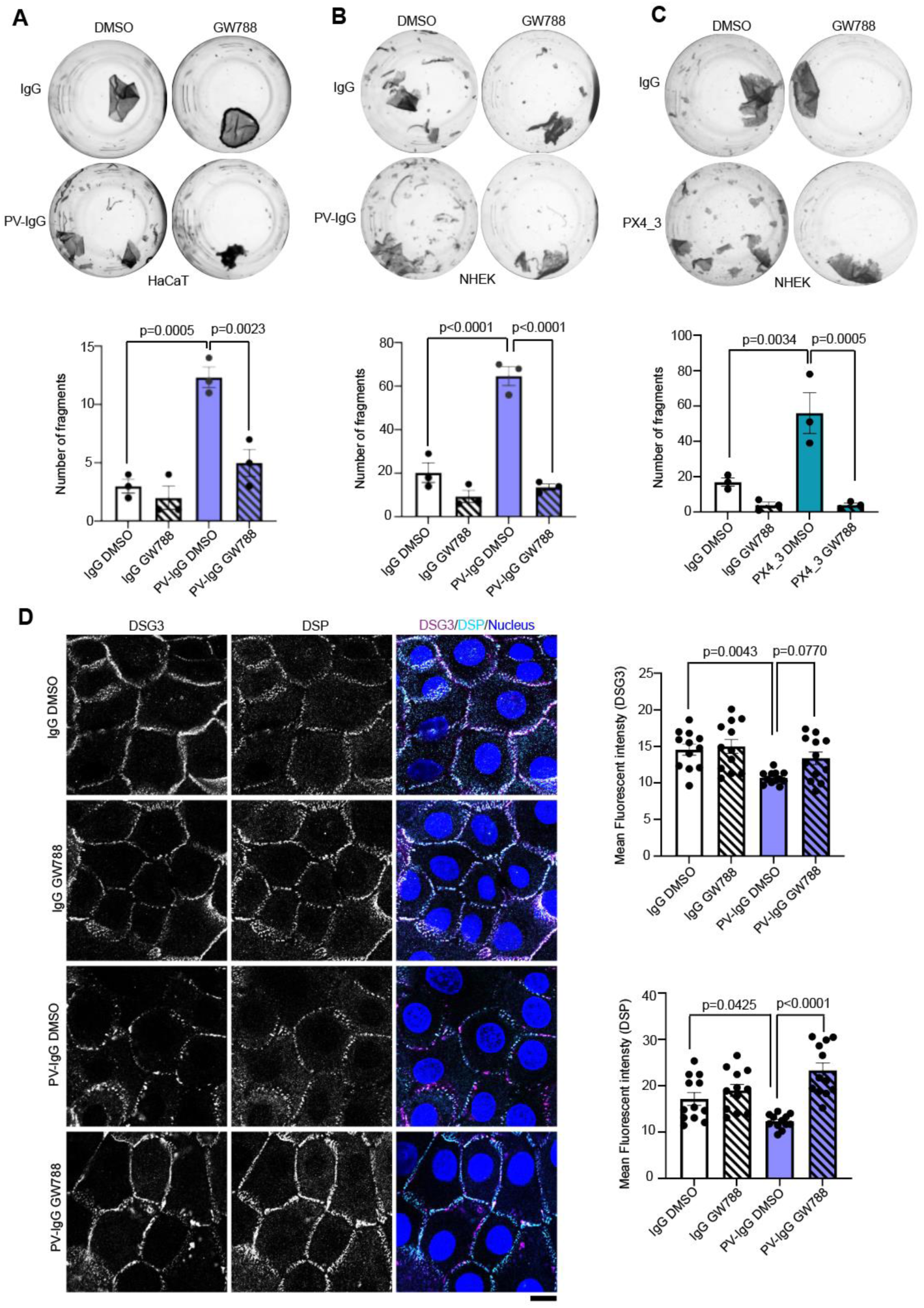
**A)** Dispase-based dissociation assay of HaCaT cells treated with IgG or PV-IgG, in combination with either DMSO or GW788 for 24 hours. Representative images and quantifications of n=3 are shown. One-way-ANOVA, Tukey’s multiple comparison used for statistical analysis. **B)** Dispase-based dissociation assay of NHEK cells treated with IgG or PV-IgG, in combination with either DMSO or GW788, for 24 hours. Representative images and quantifications of n=3 are shown. One-way-ANOVA, Tukey’s multiple comparison used for statistical analysis. **C)** Dispase-based dissociation assay of NHEK cells treated with IgG or PX4_3 in combination with either DMSO or GW788 for 24 hours. Representative images and quantifications of n=3 are shown. One-way-ANOVA, Tukey’s multiple comparison used for statistical analysis. **D)** Immunofluorescence staining of DSP and DSG3 in NHEK cells treated with IgG, PV-IgG in combination with either DMSO or GW788 for 24 hours. DAPI served to visualize nuclei. Scale bar = 10 μm. Quantification of DSG3 and DSP mean fluorescence intensity from different areas of 3 independent biological replicates are shown. One-way-ANOVA, Tukey’s multiple comparison used for statistical analysis.

We finally tested if inhibition of TGF-β influences blister formation in the skin. To do so, we applied a passive transfer human *ex-vivo* model to study the effects on blistering phenotypes of the disease. Human skin explants were injected subcutaneously with 40 ug of purified pX4_3 or IgG as a control. PX4_3 induced intraepidermal blistering after 24 hours, which was significantly reduced by concomitant injection of GW788, suggesting that TGF-β inhibition ameliorates acantholysis in human skin (**Figure 5A**). SMAD2/3 inhibition in these tissues was confirmed by injections in human *ex-vivo* skin, further supporting that PV-IgG activates TGF-β signalling.

**Figure 5.**
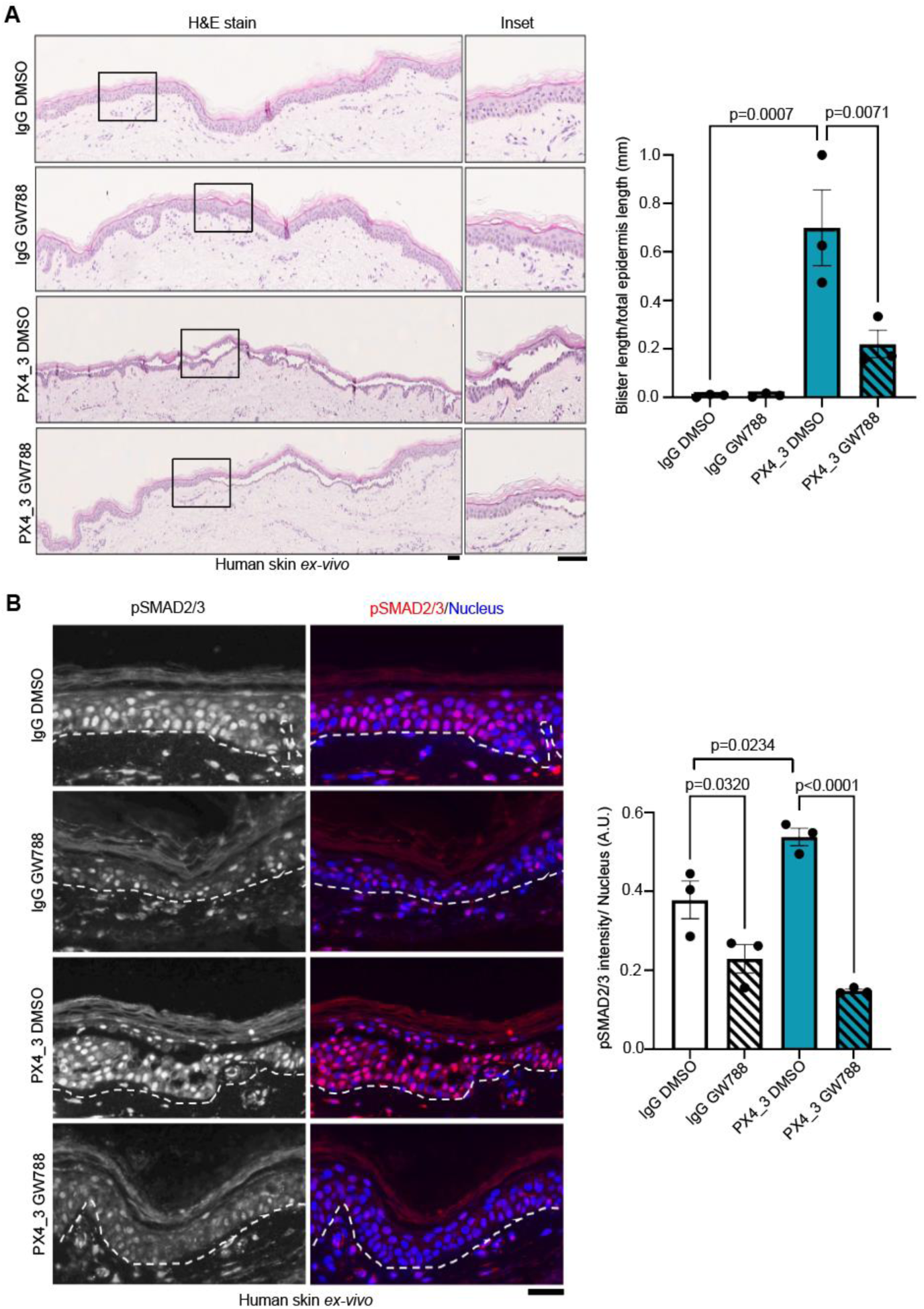
**A)** H&E staining of human skin explants injected with IgG or pX4_3 and DMSO/GW788, respectively. Scale bar = 10μm. Bar graphs represent quantification of blister length versus total epidermis length, each dot represents independent biological replicates(n=3), One-way-ANOVA, SIDAK correction used for statistical analysis. **B**) Immunofluorescence staining of human skin sections treated with Ctrl-IgG or pX4_3 and DMSO/GW788 using pSMAD2/3 (red/white) antibodies and DAPI used to stain nucleus (blue). Representative images are shown. Scale bar = 10μm. Quantification showing the mean intensity of pSMAD2/3 normalized to the respective number of nuclei within each experiment (n=3), each data point represents one biological replicate. One-way-ANOVA, SIDAK correction used for statistical analysis.

## Discussion

### SERPINB5 regulates DSP localization at the cell membrane and increases intercellular adhesive strength

From our previous study, we identified SERPINB5 as a novel interacting partner and regulatory molecule of DSP in keratinocytes^18^. In the current study, we were interested in understanding the relevance of the SERPINB5/DSP axis in a disease context with impaired adhesion. PV is an autoimmune disease with autoantibody development against the desmosomal cadherins DSG1 and DSG3, resulting in blister formation in the skin and oral mucosa. It is established that autoantibodies cause loss of cell-cell adhesion by altered turnover and stability of DSG3 and DSG1 through signalling-dependent and independent mechanisms^20^. Methods to stabilize cell-cell adhesion may represent a tailored approach to ameliorate the adverse effects of the autoantibodies from PV patients. It has been shown that overexpression of DSG3 in NHEKs prevents PV-IgG induced loss of cell-cell adhesion^21^. In a recent study, it was demonstrated that the phosphodiesterase 4 inhibitor apremilast ameliorated PV-IgG-induced blisters and cell-cell adhesion. Mechanistically, apremilast inhibited keratin retraction and promoted DSP stability through phosphorylation of plakoglobin at serine 665, which has previously been shown to be pro-adhesive ^22^ ^23^. In our study, we asked if SERPINB5-mediated regulation of DSP could, in a similar manner, improve the PV-IgG-mediated loss of cell-cell adhesion. Dispase-based cell cohesion assay showed SERPINB5-GFP overexpression in NHEK and HaCaT keratinocytes rescued PV-IgG mediated loss of cell-cell adhesion and prevented loss of DSP membrane localization under PV-IgG treatment. Interestingly, SERPINB5 did not alter the DSG3 membrane levels under these conditions, demonstrating that the effect is restricted to DSP, at least from the molecules investigated here. This is similar to the study using apremilast, where fragmentation of DSG3 staining was unaltered in response to apremilast, although loss of adhesion was blocked ^23^. It is interesting to note that modulating specific constituents of the desmosome (i.e., the transmembrane adhesion molecules, plaque proteins, and intermediate filament insertion) individually is sufficient to promote intercellular adhesion. So far it is unclear whether exclusive mechanisms are in place or whether these factors act interdependent, finally resulting in increased intercellular adhesion. Still, it is interesting to speculate that these individual targets can be exploited to precisely tune adhesion as a therapeutic principle in disease.

Our study also implicates TGF-β in the SERPINB5-dependent regulation of DSP. This is based on previous observations suggesting SERPINB5 as a negative modulator of TGF-β signalling^24^. Vice versa, TGF-β signalling has been shown to positively modulate SERPINB5, indicating the existence of a feedback loop between TGF-β activity and SERPINB5. Interestingly, TGF-β signalling was also determined to be a negative regulator of DSP localization at the cell membrane^16^. Analysis of PV patient samples showed a higher activity of SMAD2/3, a downstream effector of the TGF-β pathway. In line with patient data, treatment of keratinocytes with PV-IgG or PX4_3 enhanced the activation of SMAD2/3 and inhibition of TGF-β activation increased DSP levels at the cell membrane both under control and PV-IgG conditions. This suggests that SERPINB5 through dampening the TGF-β pathway increases DSP localization at the cell membrane, resulting in increased adhesive strength.

### Targeting TGF-β signalling as novel therapeutic approach in PV

PV impairs patient’s quality of life dramatically and can led to extreme reduction of general health status of patients. Therapeutic strategies for PV majorly include corticosteroids, steroid-sparing drugs, and B-cell suppression therapies^25^, or plasmapheresis^12,13^. Autoantibodies induce signalling pathways such as p38-MAPK and inhibition of these pathways have been tested in mouse and human skin models. More directed approaches to deplete the DSG3-specific B cells, are now being investigated, where CAAR T-cell therapy holds great promises and is under clinical trial^26^. However, due to disease heterogeneity, genetic diversity and therapy resistance, it is still important to investigate new therapeutic targets. Targeted modulation of cell-cell adhesion may serve as a supplementary measure for better management of the disease.

As we have observed TGF-β activation in PV patient samples and PV-IgG/PX4_3 treated keratinocytes, we inhibited TGF-β activation using the small molecule GW788. Importantly, inhibition of TGF-β ameliorated PV-IgG and PX4_3-mediated loss of cell-cell adhesion and blister formation in an ex-vivo passive transfer model of PV. GW788 treatment also rescued the loss of DSP from the cell-cell membrane, similar to SERPINB5 overexpression. However, PV-IgG did not modulate the expression of SERPINB5, and TGF-β activation in response to PV-IgG seems to be independent of SERPINB5 levels. The mechanisms by which PV-IgG activates TGF-β signalling need further investigation.

TGF-β activation has been associated with other skin disorders such as psoriasis and epidermolysis bullosa and squamous cell carcinoma^27–29^. TGF-β inhibition through topical application of small molecule inhibitors has been demonstrated as a therapeutic option in psoriasis, as systemic use of TGF-β inhibitors can lead to severe side effects due to pleiotropic roles of TGF-β in development and homeostasis^30^. Several ways of targeting TGF-β signalling by inhibitors of downstream effector molecules have made it possible to reduce side effects. Many of these inhibitors are in clinical trials for cancer treatments^31^. Thus, using a TGF-β inhibition strategy should be carefully designed for use in PV patients and testing the effectivity for a topical use should be considered.

In conclusion, we identify SERPINB5 as modulator of DSP membrane localization through TGF-β signalling. Further, elevated TGF-β activation was observed in PV patient samples, the inhibition of which may serve as a novel option to improve acantholysis in PV.

## Materials and Methods

### Cell culture and generation of lentiviral constructs and stable cell lines

Spontaneously immortalized HaCaT (Human adult high Calcium low Temperature) keratinocytes^32^ were cultured in a humidified atmosphere of 5% CO2 and 37°C in Dulbecco’s Modified Eagle Medium (DMEM) (Sigma-Aldrich, D6546) containing 1.8 mM Ca^2+^ and complemented with 10% fetal bovine serum (Merck, S0615), 50 U/ml penicillin (VWR, A1837.0025, D6546), 50 μg/ml streptomycin (VWR, A1852.0100) and 4 mM L-glutamine (Sigma-Aldrich, G7513). Lentiviral particles were generated according to standard procedures. HEK293T cells were transfected with lentiviral packaging vector psPAX2 (#12259, Addgene, Watertown, MA, USA), the envelope vector pMD2.G (#12260, Addgene) and the respective construct plasmid using TurboFect (Thermo Fisher Scientific, Waltham, MA, USA). 48h post transfection, virus-containing supernatant was collected and concentrated using Lenti-Concentrator (OriGene), for minimum 2h at 4°C. Cells were transduced with the respective virus particles in an equal ratio using 5 µg/mL polybrene (Sigma-Aldrich) according to the manufacturer’s instructions. 24h post transduction for HaCaT keratinocytes and 8h later for primary human keratinocytes, medium was exchanged and puromycin selection was applied. Cells were cultivated for at least one week under selective pressure, before starting with the respective experiments. Expression of the respective construct was confirmed via Western blot analysis. The cloning of constructs for SERPINB5 overexpression, sgSERPINB5, sgNT are described in the publication^24^.

### Isolation of NHEK cells

Foreskin tissue was obtained during circumcision of patients after informed consent in accordance with the local ethics committee (EKNZ; date of approval: 11.06.2018, project ID: 2018-00963). The skin samples were washed with PBS containing 300 U/mL of penicillin (#A1837, AppliChem), 300 U/mL of streptomycin sulphate (#A1852, AppliChem) and 7.5 µg/mL of amphotericin B (#A2942 Sigma-Aldrich). The dermis and epidermis were separated using 5 mg/mL Dispase II solution (#D4693, Sigma-Aldrich) in HBSS (#H8264, Sigma-Aldrich) containing 300 U/mL penicillin, 300 U/mL streptomycin sulphate and 2. 5 µg/mL amphotericin B. The detached epidermis was digested with TryplE dissociation reagent (#12605028, Gibco) containing 100 U/mL penicillin and 100 U/mL streptomycin sulfate at 37°C for 20 minutes. Following the digestion, keratinocytes were isolated by passing through a 70µm cell filter (#431751, Corning, Somerville, USA). The isolated normal human epidermal keratinocytes (NHEK) were then seeded at a density of ∼8 × 10^4^ cells/cm^2^ in EpiLife medium containing 60 µmol/L CaCl_2_ (#MEPI500CA, Gibco) and 1% human keratinocyte supplement (#S0015, Gibco), 1% Pen/Strep and 2.5 µg/mL amphotericin B. After 3 days, the medium was changed and amphotericin B was discontinued.

### Dispase-based dissociation assay

HaCaT cells were seeded in 24 well plates until 100% confluency and treated with IgG, PV-IgG or PX4_3 and drug inhibitor as indicated. The NHEK were seeded in a 24 well plate and grown to confluency, followed by addition of 1.2 mmol/L CaCl_2_ for 24 hours to induce differentiation. The cells were then washed with PBS and then incubated with 250 ul Dispase II (Sigma-Aldrich, #D4693) solution (50mg in 10 ml HBSS) for 30 minutes at 37°C, for detachment of the cell monolayer. 150 µl of HBSS (Huber Lab, #A3140) was added to the floating monolayer. The monolayer was subjected to homogeneous shear stress applied by 10x electrical pipetting with Eppendorf Xplorer 1000 (350 ul, Eppendorf, #L49475G). The fragments were documented and quantified with a stereo microscope (Olympus, #SZX2-TR30) with an attached camera (Canon, #EOS 800D).

### Western Blot

Confluent cell monolayers were lysed with SDS lysis buffer (25 mM HEPES, 2 mM EDTA, 25 mM NaF, 1% SDS, pH 7.6) supplemented with an equal volume of a protease inhibitor cocktail (cOmplete, Roche Diagnostics, Mannheim, Germany). Lysates were sonicated and the total protein amount was determined with a BCA protein assay kit (Thermo Fisher Scientific) according to the manufacturer’s instructions. The proteins were denatured by heating in Laemmle buffer, for 10min at 95°C. Membranes were blocked in Odyssey blocking buffer (Li-Cor, Lincoln, NE, USA) for 1 hour at room temperature. The following primary antibodies were diluted with odyssey blocking buffer in tris-buffered-saline containing 0.1% Tween 20 (TBS-T) (Thermo Fisher Scientific) and incubated overnight at 4°C, with rotation: rabbit SERPINB5 (MASPIN #ab182785 Abcam), mouse GAPDH mAb (clone 0411, #sc-47724 Santa Cruz), rabbit pSMAD2/3 (#AP0548 Lubioscience, Zürich, Switzerland), rabbit pSMAD1/3/5 (#ab95455 Abcam) and mouse SMAD2/3 (#sc-133098 Santa Cruz). Goat anti-mouse 800CW and goat anti-rabbit 680RD (#925-32210 and #925-68071, both Li-Cor) were used as secondary antibodies, incubated for 1hour at room temperature. Odyssey FC imaging system was used for imaging the blots and band intensity was quantified with Image Studio (both Li-Cor).

### Immunofluorescence staining and imaging

Cell were grown on coverslips till confluency and were fixed with chilled methanol (at -20°C) for 10 minutes at 4°C. Blocking was done using 3% BSA and 1% normal goat serum in PBS for 1 hour at room temperature. The coverslips were incubated overnight at 4°C with the following primary antibodies: anti-Dsg3-mAb (Invitrogen, #326300), anti-DSP-mAb (NW39, Prof. Kathleen Green laboratory). Cells were then washed three times with 1X PBS and incubated with the secondary Alexa Fluor-coupled antibodies (Fisher Scientific, A-11008, A-11004) for one hour at room temperature. DAPI (Sigma-Aldrich, D9542) was added for 10 minutes to counterstain nuclei. The coverslips were washed three more times with PBS and mounted using Prolong Diamond Antifade (Thermo Fisher Scientific, P36961). Images were taken with an HC PL APO CS2 63x/1.40 oil objective on a Stellaris 8 Falcon confocal microscope (Leica, Wetzlar, Germany).

### Passive transfer *ex-vivo* skin model

For the *ex-vivo* skin model, human skin pieces (approximately 1 cm^2^) were collected from the thigh of body donors having no history of skin disease, within 24 hours of decease. Skin at this time and after additional 24h of *ex-vivo* incubation is still viable^33^. Written informed consent was obtained from body donors for use of tissue samples for research. Skin pieces were injected superficially with 40 ug of pX4_3 or 40 ug of control IgG in PBS in combination with DMSO and GW788 (Sigma Aldrich, SML0116). These pieces were then incubated floating on DMEM (Sigma, D6546) including 10% FCS, 0.2% Glutamate and 0.5% Penicillin-Streptomycin for 24 hours. After 24 hours of incubation, a constant shear stress (10x 90° rotations with a custom-made rubber-stencil) was applied and the tissues were fixed in 4% paraformaldehyde for 24 hours. The specimens were then subjected to tissue processing, embedding in paraffin blocks and subsequent histological examination.

### Pemphigus vulgaris patient material

The usage of patient material for this study was according to the Declaration of Helsinki and was approved by the Ethics Commission of the Medical Faculty at the University of Marburg under the number 169/19. Skin punch biopsies were taken from PV patients as part of the diagnostic procedures by the Department of Dermatology, Universitätsklinikum Giessen Marburg (UKGM). The control sections were taken from excess normal skin removed during the resection of skin tumors (Supplementary Table 1). These tissues were then embedded in paraffin blocks and sectioned using a microtome. The ELISA values from PV patient sera were 1207 U/ml for DSG1 and 3906 U/ml for DSG3.

### Sex as a biological variable

The samples were used from both sexes in non-biased manner.

### Histology and immunostaining of tissue sections

Paraffin blocks were cut into 5 µm thick sections using a microtome (Thermo Fisher Scientific, HM355S). Hematoxylin and eosin (H&E) staining was carried out according to standard procedures. Briefly, the sections were stained for 5 minutes with Mayer’s hemalum solution (Sigma-Aldrich, No. 1.09249.1022), followed by washing and then dehydrated using an increasing ethanol series and counterstained with 0.5 % (w/v) eosin solution for 5 minutes. Following washing steps in ethanol and methyl salicylate, the sections were covered with DPX mounting medium (Sigma-Aldrich, #06522). For immunostaining, the sections were deparaffinized and antigen retrieval was performed in citrate buffer (10 mM citric acid monohydrate (20276.235, VWR), pH6, 0.1 % Triton) for 20 min at 95 °C. Permeabilization was done using 0.1 % Triton X-100 in PBS for 5 minutes and then blocked with 3 % bovine serum albumin/0.12 % normal goat serum in PBS for 1 hour at room temperature. The sections were incubated with the rabbit pSMAD2/3 (#AP0548 Lubioscience, Zürich, Switzerland) in PBS overnight at 4 °C. After washing, the secondary antibodies were added for 1 h, RT. DAPI (Sigma-Aldrich, D9542) was added for 10 min to stain the nuclei. Samples were embedded with Fluoromount Aqueous Mounting Medium (Sigma-Aldrich, F4680).

### Statistics and image analysis

Statistical analysis was carried out using GraphPad Prism 8 software. Data sets were first tested for normal distribution using the Shapiro-Wilk normality test. Students t-tests to compare two data sets and one-way ANOVA test for more than two data sets was performed to determine statistical significance (p<0.05). Error bars in all graphs are presented as ± SEM. A minimum of 3 independent biological replicates were used for each experiment. The figures were created with the use of Photoshop CC and Illustrator CC (Adobe, San José, CA, USA). Immunofluorescence staining of cells was analyzed with ImageJ software for quantifying DSG3, DSP and pSMAD2/3 mean intensity. Immunofluorescence staining of tissue sections were analyzed with QuPath-0.4.3. Cell nuclei were detected using DAPI, then the intensity of the protein of interest was determined for each nucleus. H&E and immune-stained samples were scanned with a slide scanner (NanoZoomerS60, Hamamatsu). Blister and total length of each H&E-stained sample was measured with NDPview.2 (Hamamatsu).

## Acknowledgements

The authors thank Dr. Diego Calabrese (Histology Core Facility), Dr. Michael Abanto and Dr. Pascal Lorentz (Microscopy Core Facility), Department of Biomedicine, University of Basel. The authors would like to thank Prof. Kathleen Green, (Northwestern University, USA) for desmoplakin antibodies and the Department of Urology (University Hospital Basel) for providing human foreskin tissue. Swiss National Science Foundation (#197764 to Prof. Dr. Spindler) and Novartis Foundation for Biomedical Research (to Prof. Dr. Spindler) for funding.

## Author Contributions

Conceptualization: VS, MR; Data acquisition: MR, AZ, HF; Data Analysis: MR, AZ, HF; Funding Acquisition: VS; Project Administration: VS, MR; Resources: VS, TC, DD, MH, ES; Supervision: VS; Writing Original Draft Preparation: MR, VS; Writing - Review and Editing: VS, MR, AZ, HF, TC, DD, MH, ES.

## Data Availability

The supporting data values from all numerical graphs are available in the XLS file.

## References

1 Nelson, W. J. The Glue that Binds Us: The Hunt for the Molecular Basis for Multicellularity. Cell 181, 495–497, doi:10.1016/j.cell.2020.03.017 (2020).

2 Waschke, J. The desmosome and pemphigus. Histochemistry and Cell Biology 130, 21–54, doi:10.1007/s00418-008-0420-0 (2008).

3 Delva, E., Tucker, D. K. & Kowalczyk, A. P. The desmosome. Cold Spring Harb Perspect Biol 1, a002543, doi:10.1101/cshperspect.a002543 (2009).

4 Green, K. J., Roth-Carter, Q., Niessen, C. M. & Nichols, S. A. Tracing the Evolutionary Origin of Desmosomes. Current Biology 30, R535–R543, doi:10.1016/j.cub.2020.03.047 (2020).

5 Broussard, J. A., Getsios, S. & Green, K. J. Desmosome regulation and signaling in disease. Cell Tissue Res 360, 501–512, doi:10.1007/s00441-015-2136-5 (2015).

6 Corrado, D., Link, M. S. & Calkins, H. Arrhythmogenic Right Ventricular Cardiomyopathy REPLY. New England Journal of Medicine 376, 1489–1490 (2017).

7 Schinner, C. et al. Defective Desmosomal Adhesion Causes Arrhythmogenic Cardiomyopathy by Involving an Integrin-alphaVbeta6/TGF-beta Signaling Cascade. Circulation 146, 1610–1626, doi:10.1161/CIRCULATIONAHA.121.057329 (2022).

8 Amagai, M., Klaus-Kovtun, V. & Stanley, J. R. Autoantibodies against a novel epithelial cadherin in pemphigus vulgaris, a disease of cell adhesion. Cell 67, 869–877, doi:10.1016/0092-8674(91)90360-b (1991).

9 Kasperkiewicz, M. et al. Pemphigus. Nat Rev Dis Primers 3, 17026, doi:10.1038/nrdp.2017.26 (2017).

10 Schmidt, E., Kasperkiewicz, M. & Joly, P. Pemphigus. Lancet 394, 882–894, doi:10.1016/S0140-6736(19)31778-7 (2019).

11 Spindler, V. et al. Mechanisms Causing Loss of Keratinocyte Cohesion in Pemphigus. J Invest Dermatol 138, 32–37, doi:10.1016/j.jid.2017.06.022 (2018).

12 Tan-Lim, R. & Bystryn, J. C. Effect of plasmapheresis therapy on circulating levels of pemphigus antibodies. J Am Acad Dermatol 22, 35–40, doi:10.1016/0190-9622(90)70004-2 (1990).

13 Ellebrecht, C. T., Maseda, D. & Payne, A. S. Pemphigus and Pemphigoid: From Disease Mechanisms to Druggable Pathways. J Invest Dermatol 142, 907–914, doi:10.1016/j.jid.2021.04.040 (2022).

14 Pemberton, P. A. et al. The tumor suppressor maspin does not undergo the stressed to relaxed transition or inhibit trypsin-like serine proteases. Evidence that maspin is not a protease inhibitory serpin. J Biol Chem 270, 15832–15837, doi:10.1074/jbc.270.26.15832 (1995).

15 Qin, L. & Zhang, M. Maspin regulates endothelial cell adhesion and migration through an integrin signaling pathway. J Biol Chem 285, 32360–32369, doi:10.1074/jbc.M110.131045 (2010).

16 Asrani, K. et al. mTORC1 loss impairs epidermal adhesion via TGF-beta/Rho kinase activation. J Clin Invest 127, 4001–4017, doi:10.1172/JCI92893 (2017).

17 Wang, S. E. et al. Convergence of p53 and transforming growth factor beta (TGFbeta) signaling on activating expression of the tumor suppressor gene maspin in mammary epithelial cells. J Biol Chem 282, 5661–5669, doi:10.1074/jbc.M608499200 (2007).

18 Rathod, M. et al. DPM1 modulates desmosomal adhesion and epidermal differentiation through SERPINB5. J Cell Biol 223, doi:10.1083/jcb.202305006 (2024).

19 Payne, A. S. et al. Genetic and functional characterization of human pemphigus vulgaris monoclonal autoantibodies isolated by phage display. J Clin Invest 115, 888–899, doi:10.1172/JCI24185 (2005).

20 Spindler, V. et al. Mechanisms Causing Loss of Keratinocyte Cohesion in Pemphigus. Journal of Investigative Dermatology 138, 32–37, doi:10.1016/j.jid.2017.06.022 (2018).

21 Jennings, J. M. et al. Desmosome disassembly in response to pemphigus vulgaris IgG occurs in distinct phases and can be reversed by expression of exogenous Dsg3. J Invest Dermatol 131, 706–718, doi:10.1038/jid.2010.389 (2011).

22 Schinner, C. et al. Adrenergic Signaling Strengthens Cardiac Myocyte Cohesion. Circ Res 120, 1305–1317, doi:10.1161/CIRCRESAHA.116.309631 (2017).

23 Sigmund, A. M. et al. Apremilast prevents blistering in human epidermis and stabilizes keratinocyte adhesion in pemphigus. Nat Commun 14, 116, doi:10.1038/s41467-022-35741-0 (2023).

24 Rathod, M., et al. DPM1 through SERPINB5 modulates desmosomal adhesion and epidermal differentiation. bioRxiv, 2022.2012.2028.522133, doi:10.1101/2022.12.28.522133 (2023).

25 Joly, P. et al. First-line rituximab combined with short-term prednisone versus prednisone alone for the treatment of pemphigus (Ritux 3): a prospective, multicentre, parallel-group, open-label randomised trial. Lancet 389, 2031–2040, doi:10.1016/S0140-6736(17)30070-3 (2017).

26 Ellebrecht, C. T. et al. Reengineering chimeric antigen receptor T cells for targeted therapy of autoimmune disease. Science 353, 179–184, doi:10.1126/science.aaf6756 (2016).

27 Han, G. et al. A role for TGFbeta signaling in the pathogenesis of psoriasis. J Invest Dermatol 130, 371–377, doi:10.1038/jid.2009.252 (2010).

28 Akasaka, E., Kleiser, S., Sengle, G., Bruckner-Tuderman, L. & Nystrom, A. Diversity of Mechanisms Underlying Latent TGF-beta Activation in Recessive Dystrophic Epidermolysis Bullosa. J Invest Dermatol 141, 1450–1460 e1459, doi:10.1016/j.jid.2020.10.024 (2021).

29 Glick, A. B. The Role of TGFbeta Signaling in Squamous Cell Cancer: Lessons from Mouse Models. J Skin Cancer 2012, 249063, doi:10.1155/2012/249063 (2012).

30 Massague, J. & Sheppard, D. TGF-beta signaling in health and disease. Cell 186, 4007–4037, doi:10.1016/j.cell.2023.07.036 (2023).

31 Kim, B. G., Malek, E., Choi, S. H., Ignatz-Hoover, J. J. & Driscoll, J. J. Novel therapies emerging in oncology to target the TGF-beta pathway. J Hematol Oncol 14, 55, doi:10.1186/s13045-021-01053-x (2021).

32 Boukamp, P. et al. Normal keratinization in a spontaneously immortalized aneuploid human keratinocyte cell line. J Cell Biol 106, 761–771, doi:10.1083/jcb.106.3.761 (1988).

33 Egu, D. T., Walter, E., Spindler, V. & Waschke, J. Inhibition of p38MAPK signalling prevents epidermal blistering and alterations of desmosome structure induced by pemphigus autoantibodies in human epidermis. Br J Dermatol 177, 1612–1618, doi:10.1111/bjd.15721 (2017).

