## Supplementary material for "SERPINB5-TGF-β signalling modulates desmoplakin membrane localization and ameliorates pemphigus vulgaris skin blistering": Supplentary file

**Supplementary Figure 1 (S1):**

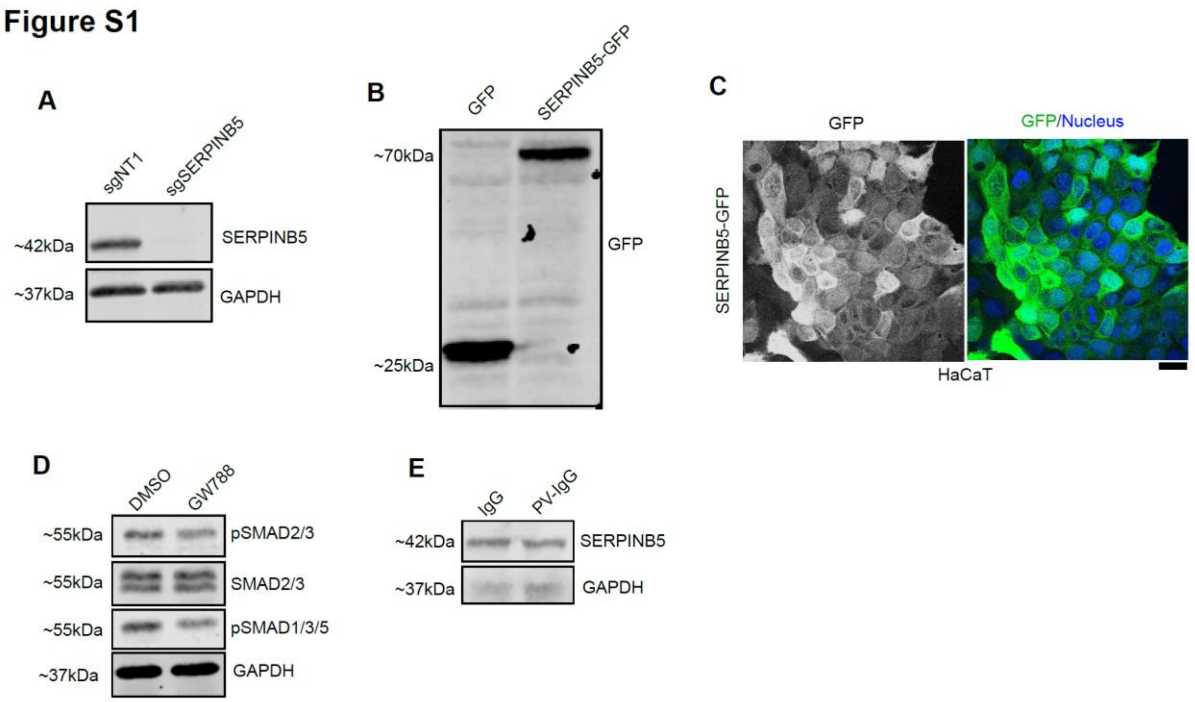


**Figure S1A)** Western blot analysis of sgNT and sgSERPINB5 HaCaT cell lysates using SERPINB5 and GAPDH antibodies to verify SERPINB5 knockdown. Representative Western blot images of 3 independent experiments are shown. **B)** Western blot image showing the expression of GFP from GFP control and SERPINB5-GFP overexpressing HaCaT cells. Representative image of 3 independent biological experiments shown. **C)** Immunofluorescence image of GFP from SERPINB5-GFP cells, to verify the expression. Green shows GFP signal and nucleus stained blue with DAPI. Scale bar = 10 $\mu$m. **D)** Western blot analysis of HaCaT cell lysates treated with DMSO, GW788388 for 24 hours using pSMAD2/3, SMAD2/3, pSMAD1/3/5 and GAPDH antibodies. Representative Western blot images and quantifications of indicated proteins (n=3) are shown. GAPDH used as loading control. **E)** Western blot image showing SERPINB5 expression in HaCaT cells treated with control IgG, PV IgG. Representative image of 3 independent biological replicates.

**Supplementary Table 1:**

| Sample | Localisation | Category | Gender (m/f) | Age (y) | anti-DSG3 | anti-DSG1 |
|---|---|---|---|---|---|---|
| 1 | skin | Ctrl | f | 84 | na | na |
| 2 | skin | Ctrl | m | 71 | na | na |
| 3 | skin | Ctrl | f | 48 | na | na |
| 4 | skin | Ctrl | f | 97 | na | na |
| 5 | skin | Ctrl | f | 48 | na | na |
| 6 | skin | Ctrl | f | 66 | na | na |
| 7 | skin | Ctrl | m | 61 | na | na |
| 8 | skin | Ctrl | m | 74 | na | na |
| 9 | skin | Ctrl | f | 86 | na | na |
| 1 | skin | Pemphigus vulgaris | m | 69 | 949 | 143 |
| 2 | skin | Pemphigus vulgaris | f | 55 | 93 | 10 |
| 3 | skin | Pemphigus vulgaris | f | 83 | 141 | 861 |
| 4 | skin | Pemphigus vulgaris | f | 65 | 935 | 737 |
| 5 | skin | Pemphigus vulgaris | f | 65 | 178 | 3 |
| 6 | skin | Pemphigus vulgaris | m | 86 | 118 | <20 |
| 7 | skin | Pemphigus vulgaris | m | 86 | 118 | <20 |

**Table 1:** Table showing the patient information from control and PV. The anti-DSG3 and anti-DSG1 values indicated are titre values from ELISA assay (U/ml). Na = not detected. m/f indicates male/female respectively.
